# Species-specific, age-varying plant traits affect herbivore growth and survival

**DOI:** 10.1101/2020.01.14.906859

**Authors:** Louie H. Yang, Meredith L. Cenzer, Laura J. Morgan, Griffin W. Hall

**Author notes:** Department of Ecology and Evolution, University of Chicago, Chicago, IL 60637 USA.

## Abstract

Seasonal windows of opportunity represent intervals of time within a year during which organisms have improved prospects of achieving life history aims such as growth or reproduction, and may be commonly structured by temporal variation in abiotic factors, bottom-up factors, and top-down factors. Although seasonal windows of opportunity are likely to be common, few studies have examined the factors that structure seasonal windows of opportunity in time. Here, we experimentally manipulated host plant age in two milkweed species (*Asclepias fascicularis* and *Asclepias speciosa*) in order to investigate the role of plant species-specific and plant age-varying traits on the survival and growth of monarch caterpillars (*Danaus plexippus*). We show that the two plant species showed diverging trajectories of defense traits with increasing age. These species-specific and age-varying host plant traits significantly affected the growth and survival of monarch caterpillars through both resource quality- and resource quantity-based constraints. The effects of plant age on monarch developmental success were comparable to and sometimes larger than those of plant species identity. We conclude that species-specific and age-varying plant traits are likely to be important factors with the potential to structure seasonal windows of opportunity for monarch development, and examine the implications of these findings for both broader patterns in the ontogeny of plant defense traits and the specific ecology of milkweed-monarch interactions in a changing world.

## Introduction

Seasonal windows of opportunity are intervals of time within a year during which organisms have improved prospects of achieving life history aims such as growth or reproduction (Yang and Cenzer 2019). Seasonal windows of opportunity are likely to occur in a wide range of systems (e.g., Yang and Rudolf 2010, Anderson et al. 2012, Wright et al. 2013, Carter et al. 2018, Farzan and Yang 2018, Yang and Cenzer 2019), resulting from commonplace temporal variation in biotic and abiotic factors. However, while phenology examines the *realized* seasonal timing of an organism’s life history, seasonal windows of opportunity represent transient periods of time with the *potential* for improved developmental or fitness outcomes. Because underlying windows of opportunity may not always be reflected in observed phenology, experimental manipulations provide a particularly useful approach for identifying seasonal windows of opportunity (Yang and Rudolf 2010). Despite this, relatively few studies have experimentally identified seasonal window of opportunity in nature (but see Van Asch et al. 2007, Liu et al. 2011, Rafferty and Ives 2011, Warren et al. 2011, Kharouba et al. 2015, Farzan and Yang 2018, Yang and Cenzer 2019), and even fewer have experimentally examined the specific factors that define these windows of opportunity in time.

Seasonal windows of opportunity are defined by the co-occurrence of factors that, in combination, have a positive effect on growth or reproduction. Broadly, many seasonal windows of opportunity are likely to be structured by temporal variation in abiotic factors, bottom-up factors, and top-down factors (Yang and Cenzer 2019). When the combined effects of these factors present adverse conditions, they constrain the seasonal timing of development. When the combined effects of these factors are favorable, they create seasonal windows of opportunity.

However, separating and evaluating the role of specific factors in structuring seasonal windows of opportunity is challenging due to the multiple correlated factors that often change simultaneously across a seasonal timescale.

The interactions between herbivores, their host plants, and their surrounding community provide unique opportunities to examine seasonal windows of opportunities. For herbivores, these windows of opportunity are likely to be structured by a variety of seasonally varying factors, including climatic conditions, natural enemy communities and plant traits. Questions about the ontogeny of plant defense traits have received particular attention as ecologists have sought to understand the specific mechanisms (Barton 2013, 2016, Quintero et al. 2013) and general patterns (Boege and Marquis 2005, Barton and Koricheva 2010, Barton and Boege 2017) that explain how plant-herbivore interactions change across development. Broadly, these studies document a diversity of ontogenetic trajectories (including both increasing and declining trends) in a wide range of plant defense traits (including both tolerance and chemical, physical, and indirect resistance traits). While specific patterns of change differ with both plant and herbivore identity (Barton and Koricheva 2010), the observation of significant ontogenetic changes in plant defense traits is both general and robust (Barton and Koricheva 2010, Barton and Boege 2017). In addition, plant phenology has recently been suggested as a key factor that could unify the hypothesis that herbivores generally prefer and perform better on vigorously growing plants (i.e., the *plant vigor hypothesis*, Price 1991) and the hypothesis that herbivore outbreaks are more likely on stressed plants (i.e., the *plant stress hypothesis*, White 1974); phenological changes in plant traits can change the quality of plant resources in ways that are consistent with both hypotheses (White 2009, Che-Castaldo et al. 2019). However, while seasonal changes in plant defense traits are likely to be a common consequence of plant ontogenetic trajectories in many systems, few studies have examined the ecological consequences of these temporally variable plant defense traits for the developmental prospects of herbivores.

Here, we present an experiment designed to isolate and examine the role of plant traits in constraining seasonal windows of opportunity for larval monarchs (*Danaus plexippus*) feeding on two milkweed host plants (*Asclepias fascicularis* and *Asclepias speciosa*). While previous studies have identified seasonal windows of opportunity in the interactions between milkweed host plants and their monarch caterpillar herbivores (Yang and Cenzer 2019), more specific experiments are necessary to identify the factors that structure these windows of opportunity in time. In this experiment, we isolated the species-specific effects of age-varying plant traits on the developmental prospects of monarch caterpillars by presenting plants of two milkweed species and three age classes synchronously to a single cohort of monarch caterpillars. This design aimed to control for the effects of seasonally variable abiotic conditions and natural enemy communities while isolating the effects of species-specific and age-varying plant traits. The key questions we address in this study are: a) How do plant traits, including measures of both size (i.e., resource *quantity*) and defensive traits (i.e., resource *quality*), change with plant age in two species of milkweed host plants? b) How do these species-specific and age-varying changes in plant traits affect the growth and survival of larval monarchs?

## Methods

### Plant establishment

We started three cohorts of narrow-leaved milkweed (*A. fascicularis*) and showy milkweed (*A. speciosa*) from seed on April 8, May 7 and June 8, 2014. These two milkweed species are native to the California Central Valley, and the seeds used in this study were propagated from local source populations (Hedgerow Farms, Winters, CA, USA). Each cohort of seeds was started directly into 2.5 L containers filled with potting soil (1:1:1 ratio of sand, compost and peat moss by volume with 1.78 kg/m^3^ dolomite), which were irrigated and fertilized (electrical conductivity, EC = 1.5-1.6 mS cm^-1^) via drip emitters as necessary to prevent water and nutrient limitation. Plants from each cohort were randomly interspersed in a single greenhouse (approximately 20-35° C) at the University of California, Davis Orchard Park Research Greenhouse Facility (38.543129° N, 121.763425° W) with individual plants spaced on open grate wire benches to prevent contact between the leaves of neighboring plants. These three cohorts were started approximately 4 weeks apart to yield three distinct age classes of milkweed (25-day, 57-day and 86-day-old plants, hereafter, the *4, 8* and *12-week* cohorts) for each species (*N*=18 plants of each species in each age class, *N*=108 plants total) at the start of the experiment.

### Measuring plant traits

We measured the size (total stem length, total leaf count, total stem cross-sectional area and total leaf area) and defensive traits (mean latex exudation and trichome density) of each plant at the start of the experiment (July 3, 2014). All plants were actively growing at the start of the experiment, and two of the 12-week-old plants had begun developing flowers (reflecting *seedling, vegetative juvenile* and *juvenile-mature transition stages, sensu* Barton and Koricheva 2010). In the context of this experiment, plant age provides a proxy for both plant phenology and ontogeny; i.e., older plants represent plants that are more phenologically advanced and developmentally mature. Total stem length was measured as the product of the total stem count (all stems > 5 cm), and the mean stem length (averaged from a subsample of up to 10 stems > 5 cm in length). Total leaf counts included all fully expanded leaves on each plant. Total stem cross-sectional area is the cumulative cross-sectional area of stems, calculated from the total stem count (all stems >5 cm) and the mean stem diameter measured from a subsample of up to 10 stems >5 cm in length. Total leaf area was estimated as the product of the total leaf count and the mean area per leaf for each plant species × plant age combination. The mean area per leaf was estimated as the area of an ellipse using measurements of the length and width of *N*=5 fully expanded leaves randomly selected from each group. Latex exudation was measured as the mean dry mass of latex collected on pre-weighed filter paper discs after cutting 5 mm from the distal tip of two fully expanded upper leaves, following Agrawal (2005). Trichome density was assessed from the upper surface of 3 mm diameter leaf discs punched from fully expanded apical leaves using digital analysis of magnified images to determine the proportion of the leaf area obscured by trichomes based on manual color thresholding (Abramoff et al. 2004).

### Monarch introduction and monitoring

A single monarch egg was introduced to each plant on July 3, 2014 (experimental day 0). In order to minimize direct handling of the eggs, we punched 6.4 mm leaf discs from oviposition host plants with single monarch eggs attached, and attached them to the apical leaves on their experimental host plants with a drop of milkweed latex. Monarch eggs were obtained from a large, local insectary population (Utterback Farms, Woodland, CA, USA) which was re-established from local monarch genotypes each year, maintained in large greenhouses, regularly supplemented with new adults to maintain genetic diversity, and had been previously assessed for parasites and pathogens (H.K. Kaya, *pers. comm.*). All monarch eggs in this experiment were selected haphazardly from a single oviposition time-restricted cohort to minimize variation in hatch timing. Each monarch egg was checked 24 h after its initial introduction (experimental day 1) to assess hatch rate and larval length. Afterwards, we re-measured caterpillars every 2-3 days until they died or left the plant (*N*=1034 observations). All larvae were measured to the nearest 0.1 mm using dial calipers; eggs were assumed to have a length of zero. Larval mass was estimated from a power law regression of caterpillar length and mass, parameterized from a dataset describing 73 unmanipulated caterpillars measured in 2014 (mass=0.0223 * length + 2.9816, *R^2^*=0.97). During each observation, we also visually estimated the proportion of leaf area that was removed due to herbivory (hereafter, *percent damaged*). Caterpillars were intentionally not bagged or constrained at any point in this experiment so that we could assess when caterpillars left their host plants (in terms of caterpillar age, caterpillar size, and host plant herbivory). Caterpillars that left their host plant below a minimum threshold size for pupation (35 mm length, or 895 mg) were assumed to have been unable to complete their larval development on a single host plant; in the context of a single plant patch, we considered these to be “dead” in our survival analyses. Caterpillars that left their host plant after attaining this threshold size were considered to be seeking pupation sites, and were considered to be right-censored in survival analyses. The threshold size for pupation (895 mg or 35 mm) was determined by assessing the larval size attained by all pupating caterpillars in previous field experiments, and among 248 caterpillars reared in the laboratory in 2014 and 2015 (Yang and Cenzer 2019). In 2.8% (*N*=29) of observations, we observed a second non-focal caterpillar that had moved onto an experimental plant; in the majority of these cases, we were able to unambiguously identify the focal caterpillar and remove the non-focal caterpillar. In three instances (0.3% of observations), the identity of the focal caterpillar could not be determined; although the qualitative conclusions of this study were unaffected by the inclusion or exclusion of these plants, we removed all observations from these three plants for the analyses presented here.

### Analyses of plant traits

We analyzed plant traits (total stem length, total stem cross-sectional area, total leaf area, mean latex exudation and trichome density) using linear models with likelihood ratio tests to assess the significance of plant species, plant age and their interaction as explanatory categorical factors (R Core Team 2018). These analyses allowed us to examine how plant traits changed with age in each milkweed species.

### Survival analyses

We analyzed the survival of monarchs for each plant species and age cohort to generate species- and age-specific Kaplan-Meier survivorship curves (Therneau and Grambsch 2000, Therneau 2015, Kassambara and Kosinski 2019). We compared curves using a log-rank test procedure for right-censored data (Harrington and Fleming 1982) implemented in the *survdiff* function in the *survival* package in R (Therneau 2015). We quantified the overall daily survivorship rates for each group of interest using the slope coefficient of a log-linear regression of survival rates over time, with visual inspection to confirm model fit assumptions. In addition, we used a Cox proportional hazards model in order to combine plant species and plant age effects into a single survival model (using the *coxph* function in the *survival* package, Therneau 2015) and estimate the proportional hazard ratios associated with the specific levels of each factor (using the *ggforest* function in the *survminer* package, Kassambara and Kosinski 2019).

### Estimation of larval growth rates

We estimated overall larval growth rates as the slope of the log-linear fit of experimental day vs. log(mass) for each individual caterpillar; i.e., as a relative growth rate. In order to estimate the slope of a log-linear regression in a dataset that included zero values, we added a small constant equal to the minimum observed mass across the dataset to all mass data in the log-linear analysis. We used a log-linear fit of mass (as opposed to length) data because visual inspection indicated that caterpillar masses show a more log-linear (i.e. exponential) pattern of increase over time, although these two metrics of monarch size yield qualitatively identical results. To avoid inaccurate overall slope estimates resulting from insufficient data, we excluded caterpillars that died before reaching 10 mm length.

In addition, we also estimated overall larval growth rates as the mass of caterpillars on experimental day 8; i.e., as the absolute growth rate. When assessing caterpillar size attained over this interval, all caterpillars that did not survive to the end of that interval were necessarily excluded. We chose day 8 for these growth rate estimates in order to achieve a balance between maximizing the length of time considered, and minimizing the number of caterpillars excluded.

For simplicity, we primarily present relative growth rates based on the slope of the log-linear regression here because this estimate is informed by more observations for each summary growth rate, and because this approach can be more easily generalized to examine a range of interval-specific growth rates. Because both of these overall growth rate estimates are measured relative to size on day 0, they are mathematically similar and yield qualitatively similar results; in addition, although they use different criteria for data exclusion, they both summarize the growth rates of a similar number of caterpillars (*N*=74 for the log-linear approach, and *N*=71 for the size on day 8 approach). For completeness, the analysis of absolute growth rates is presented in Appendix S1.

We also estimated the interval-specific relative growth rates of caterpillars using log-linear regression on two timescales: a) for all possible intervals; i.e., between all available adjacent experimental days (0, 1, 4, 6, 8, 11, 13, 15, and 18) and b) comparing early (between days 0 and 1) and late (between days 1 and 11) growth rates.

### Analyses of plant species and plant age effect sizes on larval growth rates

We calculated the size of the plant species effect for each cohort as the fixed effect coefficient of the plant species factor in a linear model with the overall relative growth rate as the response variable. This effect size metric describes the expected proportional change in the relative growth rate for caterpillars reared on showy milkweed relative to narrow-leaved milkweed. An effect sizes of would zero indicate that caterpillars showed similar relative growth rates on narrow-leaved and showy milkweed; negative effect sizes indicate that growth rates were slower on showy milkweed than on narrow-leaved milkweed. For example, an effect size of −0.05 for a given cohort would indicate that the caterpillars in that cohort showed relative growth rates that are 5% lower on showy milkweed than on narrow-leaved milkweed.

We also calculated the size of the plant age effect for each available experimental day (0, 1, 4, 6, 8, 11, 13, and 15) and plant species combination using the fixed effect coefficient of the plant age explanatory factor in a linear model with log-transformed mass as the response variable. This effect size metric describes the effect of plant age on the overall relative growth rate of caterpillars on each plant species for each day of the experiment in units of proportional change in mass per week. In this analysis, an effect size of zero would indicate that caterpillar mass was uncorrelated with plant age on a given experimental day; negative effect sizes indicate that plant age was negatively correlated with caterpillar mass. For example, an effect size of −0.05 in this analysis would indicate that the expected mass of surviving caterpillars on a given experimental day, developing on a given host plant species was reduced by 5% for each week of increasing host plant age.

### Analyses of maximum larval size attained

We analyzed the maximum larval size attained using linear models and likelihood ratio tests to evaluate the significance of plant species, plant age and their interaction effects as explanatory categorical factors (R Core Team 2018). Maximum larval size provides an integrated measurement of larval developmental success including aspects of both growth and survival.

### Analyses of plant damage

We analyzed the maximum percent damaged using linear models and significance tests with plant species, plant age and their interaction as explanatory categorical factors (R Core Team 2018), as in the analysis of maximum larval size. Maximum percent damaged indicates the maximum level of herbivory before the caterpillar died or left the plant.

## Results

### Plant traits varied with plant species and age

The size and defensive traits of both milkweed species changed over time in species-specific ways. Across all cohorts, narrow-leaved milkweed showed total stem lengths that were 3.1 times greater than those of showy milkweed (*plant species: F_1,106_*=76.7, *p*<0.0001, Fig. 1a). While both species increased their total stem length across the three cohorts (*plant age: F_1,106_*=128.5, *p*<0.0001), total stem length increased more quickly in narrow-leaved milkweed than in showy milkweed (*plant species × plant age: F_1,105_*=117.3, *p*<0.0001), reflecting differences in the architecture of these two species. In 4-week-old plants, the mean total stem length of narrow-leaved milkweeds was only 1.2 times that of showy milkweed (12.5 vs. 10.4 cm), but this difference increased to 3.3 times (44.9 vs. 13.7 cm) in 9-week-old plants, and to 3.6 times in 12-week-old plants (116.3 vs. 31.9 cm). Total leaf count showed a similar pattern (Fig. 1b). The total cross-sectional stem area was also greater in narrow-leaved milkweed overall (*plant species*: *F_1,106_*=14.6, *p*=0.0002, Fig. 1c), increased with plant age (*plant age: F_1,106_*=180.4, *p*<0.0001); and increased more in narrow-leaved milkweed relative to showy milkweed (*plant species × plant age: F_1,105_*=4.2, *p*=0.041), though this weaker interaction effect suggests that this metric of plant size did not continue to diverge over plant ontogeny (Fig 1c). By comparison, total leaf area increased with plant age (*plant age: F_1,106_*=285.3, *p*<0.0001, Fig. 1d), but did not differ between species overall (*plant species*: *F_1,106_*=0.028, *p*=0.867, Fig. 1d); while narrow-leaved milkweed showed an accelerating trajectory of increasing leaf area with age, showy milkweed showed a decelerating trajectory of increasing leaf area with age (*plant species × plant age: F_1,105_*=8.6, *p*=0.0041, Fig. 1d).

**Figure 1.**
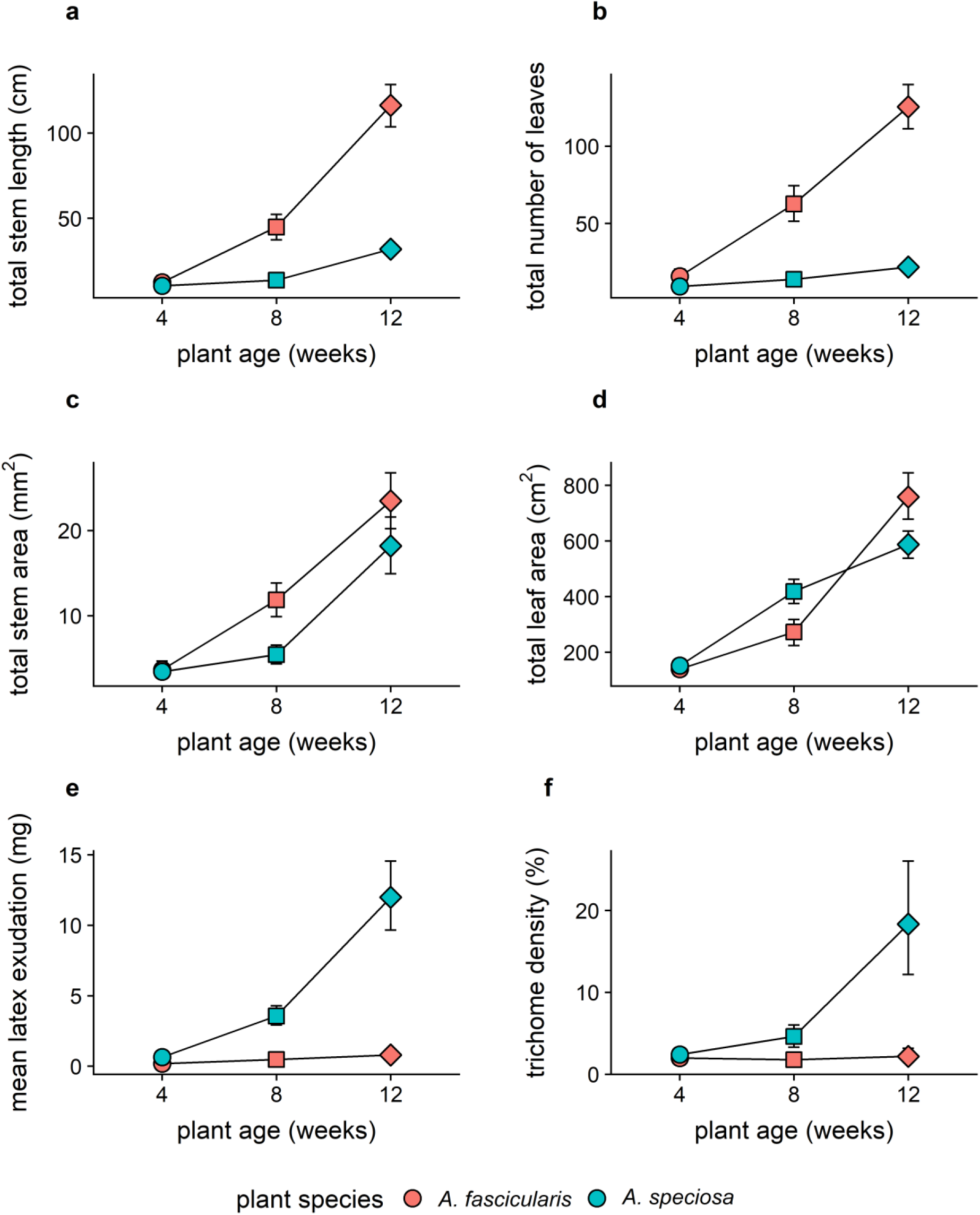
Plant traits a) mean total stem length, b) mean total leaf count, c) total stem cross-sectional area, d) total leaf area, e) mean latex exudation, and d) mean trichome density changed over plant ontogeny and differed between plant species. Color represents plant species, and point shape represents plant age. Error bars represent 95% confidence intervals.

In contrast, both defense traits showed a significant diverging pattern with plant age (Fig 1e and 1f). Overall, mean latex exudation was 11 times greater in showy milkweed compared to narrow-leaved milkweed (*plant species: F_1,106_*=57.3, *p*<0.0001, Fig. 1e), and the mass of exuded latex increased with plant age for both species (*plant age: F_1,106_*=55.8, *p*<0.0001, Fig. 1e). However, the pattern of increased latex exudation with plant age differed strongly by plant species (*plant species × plant age: F_1,105_*=77.6, *p*<0.0001, Fig. 1e); while the mean mass of exuded latex increased more than four-fold between 4 and 12 week-old narrow-leaved milkweeds (0.19 mg to 0.80 mg), it increased by almost 19 times between 4 and 12 week-old showy milkweeds (0.64 mg to 12.00 mg). Among 4-week-old plants, showy milkweed exuded 3.4 times more latex than narrow-leaved milkweed (0.64 vs. 0.19 mg); among 12-week-old plants, showy milkweed exuded 14.9 times more latex than narrow-leaved milkweed (12.00 vs. 0.80 mg). Trichome densities showed a similar pattern; overall, trichomes were 4.2 times denser on showy milkweed compared with narrow-leaved milkweed (*plant species*: *F_1,106_*=19.2, *p*<0.0001, Fig. 1f), and plants showed generally increasing mean trichome densities with plant age across both species (2.2% among 4-week-old plants to 10.2% among 12-week-old plants, *plant age: F_1,106_*=19.5, *p*<0.0001, Fig. 1f). Trichome densities increased faster on showy milkweed than on narrow-leaved milkweed (*plant species × plant age: F_1,105_*=22.3, *p*<0.0001, Fig. 1f).

Plant age explained more of the observed variation in total stem length, total stem cross-sectional area and total leaf area than plant species (*ΔR^2^*=0.41 vs *ΔR^2^*=0.25 for total stem length, *ΔR^2^*=0.60 vs *ΔR^2^*=0.05 for total stem cross-sectional area, *ΔR^2^*=0.73 vs *ΔR^2^*=0.0001 for total stem length). The variance explained by plant age and plant species was comparable for total leaf count (*ΔR^2^*=0.31 for plant age vs. *ΔR^2^*=0.35 for plant species), latex exudation (*ΔR^2^*=0.26 for plant age vs *ΔR^2^*=0.26 plant species) and trichome density (*ΔR^2^*=0.14 for plant age vs *ΔR^2^*=0.13 plant species).

### Plant species and plant age effects on larval survival

Across all cohorts, the survival curves of monarch larvae differed on narrow-leaved and showy milkweed (*χ^2^_1_*=4.8, *p*=0.028), with caterpillars on narrow-leaved milkweed showing 10.4% higher daily survival rates (91.6% vs 82.9%, Fig. 2). This result is consistent with the increased hazard ratio (1.59, 95% CI 1.04-2.5, *p*=0.034) observed on showy milkweed relative to narrow-leaved milkweed (Fig. S1). This effect of plant species on survival became stronger with plant age; while the survival curves of caterpillars on both host plant species are largely overlapping for 4-week-old plants (*χ^2^_1_*=0, *p*=0.99, Fig. 2a), they are more different on 8- and 12-week-old plants (8-week-old plants: *χ^2^_1_* =2.9, *p*=0.089, Fig. 2b; 12-week-old plants: *χ^2^_1_* =2.9, *p*=0.086, Fig. 2c). For example, caterpillars showed 2.4% greater daily survival rate on showy milkweed among 4-week-old plants (Fig. 2a), but showed 10.1% and 8.4% greater daily survival on narrow-leaved milkweed in weeks 8 and 12, respectively (Fig. 2b and 2c). We did not observe a statistically significant overall effect of plant age on the survival curves of larvae developing on either host plant species using log-rank tests (narrow-leaved milkweed, *χ^2^_2_*=2.8, *p*=0.247; showy milkweed, *χ^2^_2_*=0.8, *p*=0.684), although a comparison between the youngest and oldest plant age groups suggested a stronger pattern of lower survival on younger plants of narrow-leaved milkweed (*χ^2^_1_*=2.9, *p*=0.0885) compared to showy milkweed (*χ^2^_1_*=0.4, *p*=0.523). However, we did observe a trend towards reduced survival on younger plants across both species, which was consistent with the estimated hazard ratios for 8-week-old plants (0.93, 95% CI 0.57-1.5, *p*=0.792) and 12-week-old plants (0.70, 95% CI 0.41-1.2, *p*=0.195) relative to 4-week-old plants (Fig. S1). Overall, caterpillars on both host plants species showed the lowest daily survival rates on the youngest host plants (Fig. 2 and S2, 79.5% on narrow-leaved milkweed, 81.5% on showy milkweed), with increasing daily survival rates on older plants (8-week-old plants: 92.8% on narrow-leaved milkweed, 84.3% on showy milkweed; 12-week-old plants: 96.6% on narrow-leaved milkweed, 89.1% on showy milkweed).

**Figure 2.**
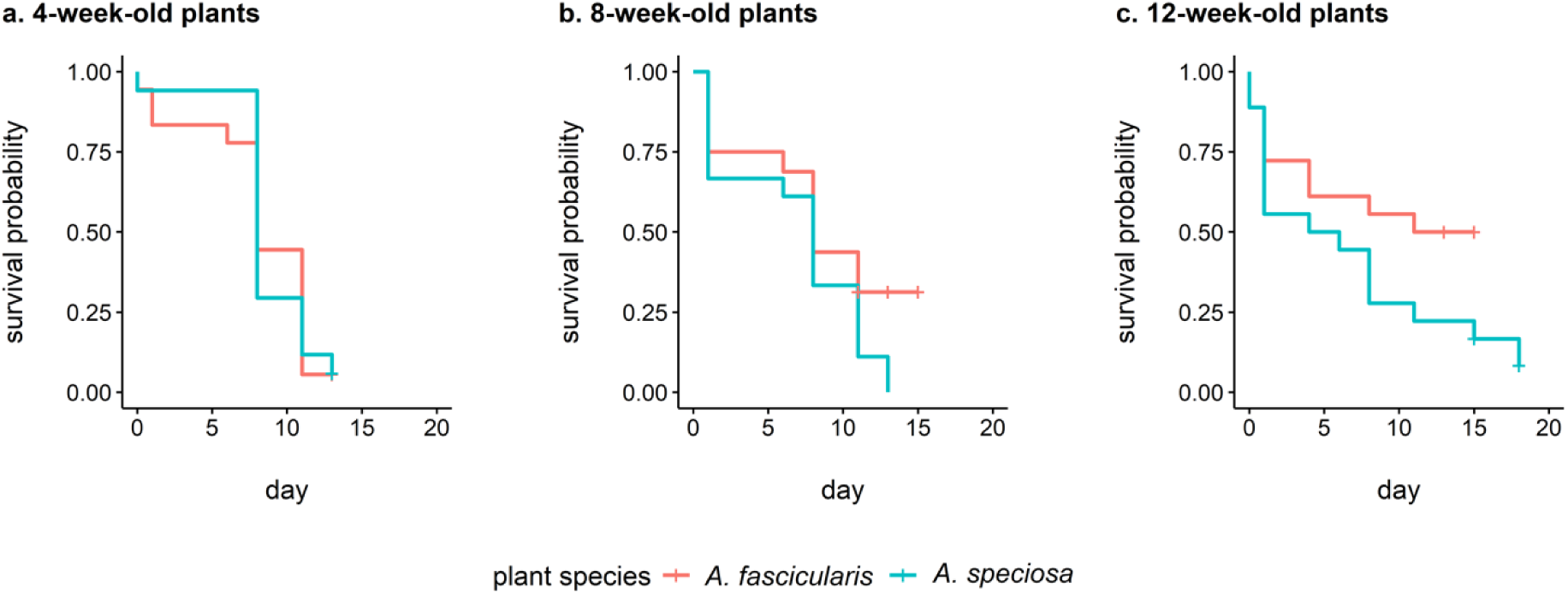
Survival of larval monarchs on a) 4-week-old, b) 8-week-old and c) 12-week-old plants. Tick marks on the survivorship curve indicate pupation. Color represents plant species.

### Plant species and plant age effects on larval growth rates

Across all host plant cohorts, larval growth was 5.7% higher on narrow-leaved milkweed than on showy milkweed (0.79 mg/mg/day vs. 0.74 mg/mg/day; *plant species*, *F_1,71_*=4.0, *p*=0.049, Fig. 3-5), with no significant differences in the effects of plant age on larval growth across species (*plant species × plant age: F_2,70_*=1.53, *p*=0.22). However, developing on showy milkweed (instead of narrow-leaved milkweed) had negative effects on relative growth rate that were 4.2 times greater in 12-week-old plants compared with 4-week-old plants (Fig. 6a; 4-week-old plants, −0.027 mg/mg/day; 8-week-old plants, −0.016 mg/mg/day; 12-week-old plants, −0.114 mg/mg/day, Fig. 6a). This result suggests that species-specific differences in plant traits on monarch growth are stronger in older plants than in younger plants. Overall, plant age explained 5 times more variation in overall larval growth rate than plant species (*ΔR^2^*=0.207 for plant age*, ΔR^2^*=0.043 for plant species).

**Figure 3.**
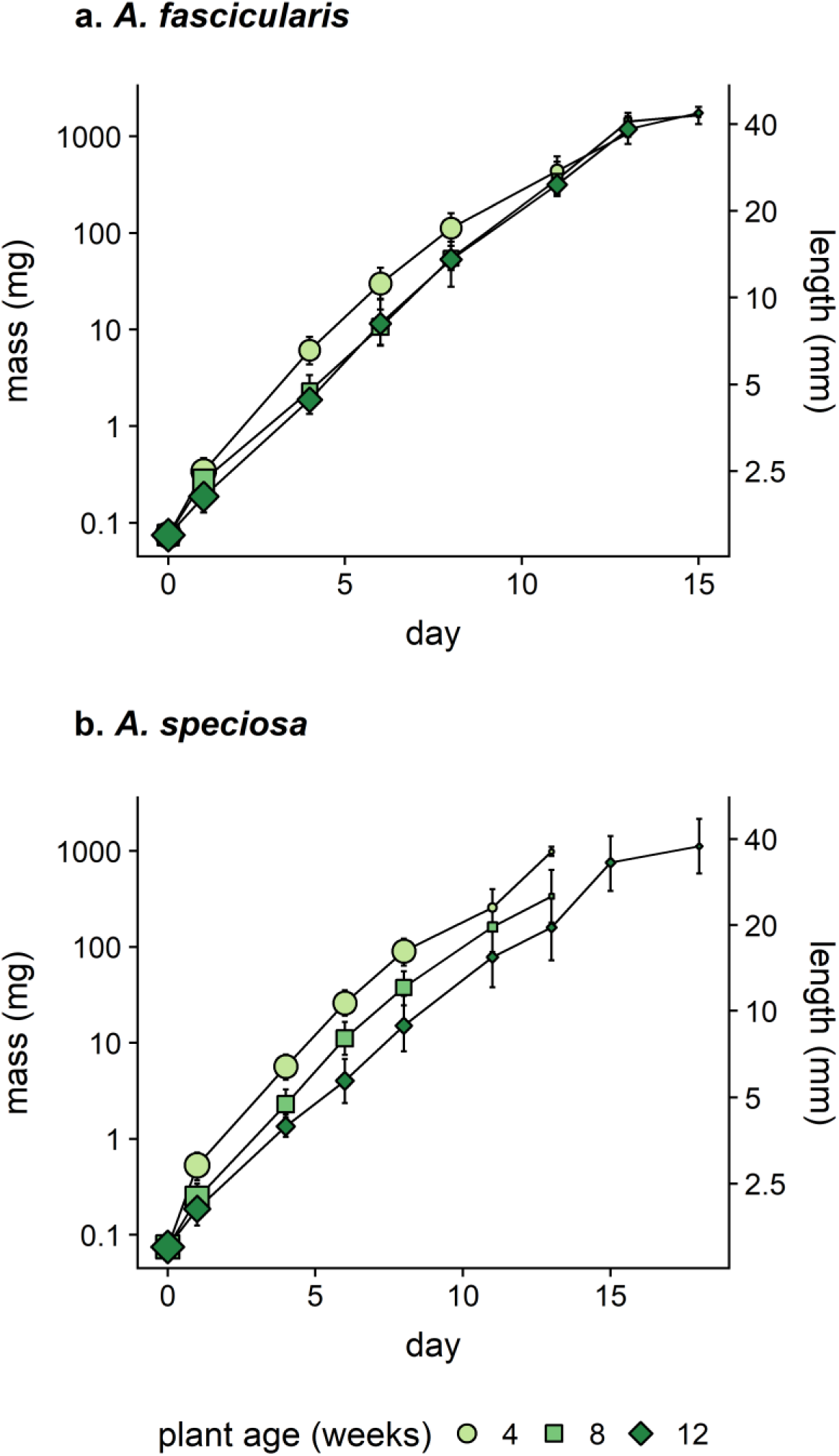
Mean surviving larval size over time for caterpillars developing on a) narrow-leaved milkweed and b) showy milkweed host plants, plotted on a log scale. The minimum observed mass value (0.75 mg) was added to each observation to allow the plotting of observed zero values. Host plant age affects larval size on both plant species throughout the experiment. Point area reflects the size of the surviving population, color and point shape represent plant age. Error bars represent 95% confidence intervals.

**Figure 4.**
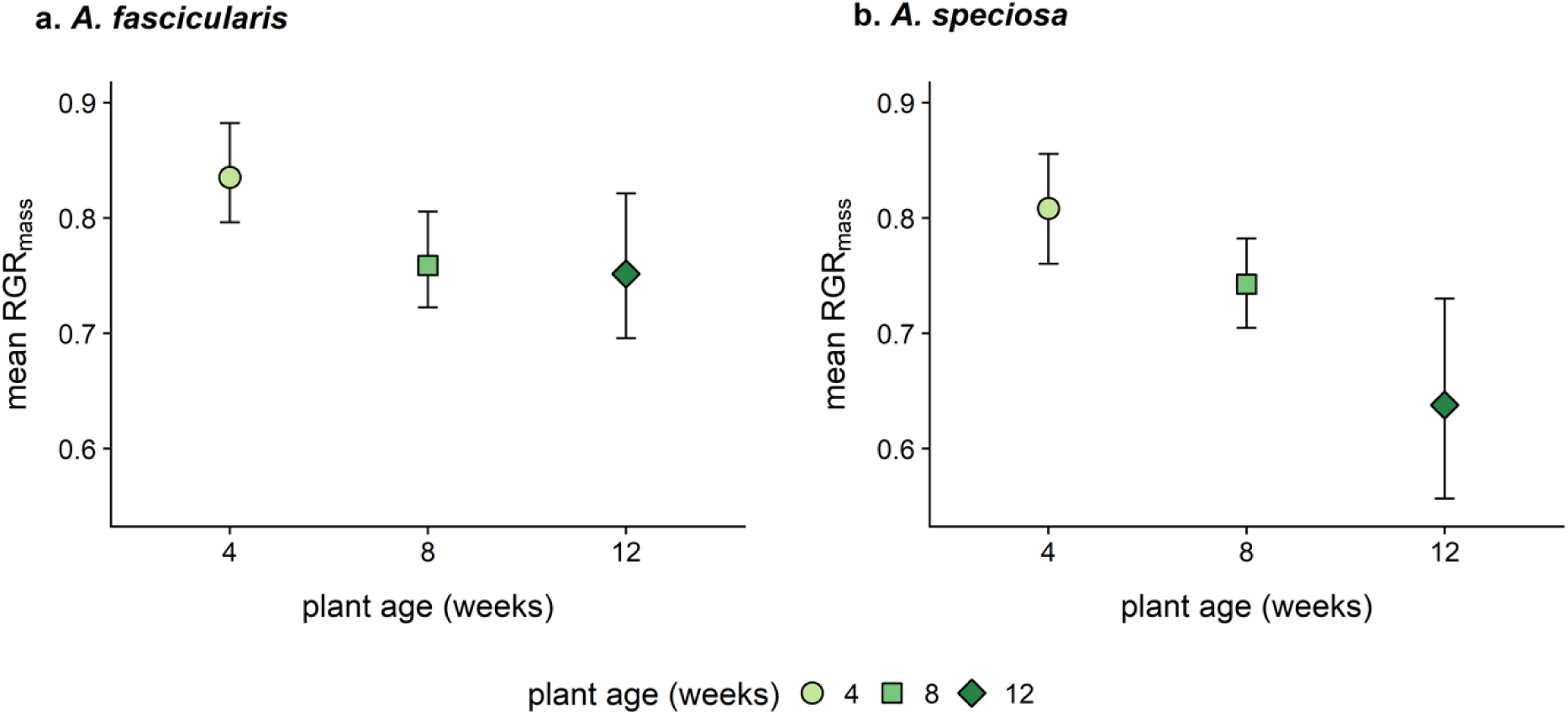
Overall mean relative growth rates for caterpillars developing on each plant age cohort of a) narrow-leaved milkweed and b) showy milkweed. This figures shows an effect of plant age on the overall (lifetime) growth rates of caterpillars. Point color and point shape represent plant age. Error bars represent 95% confidence intervals.

**Figure 5.**
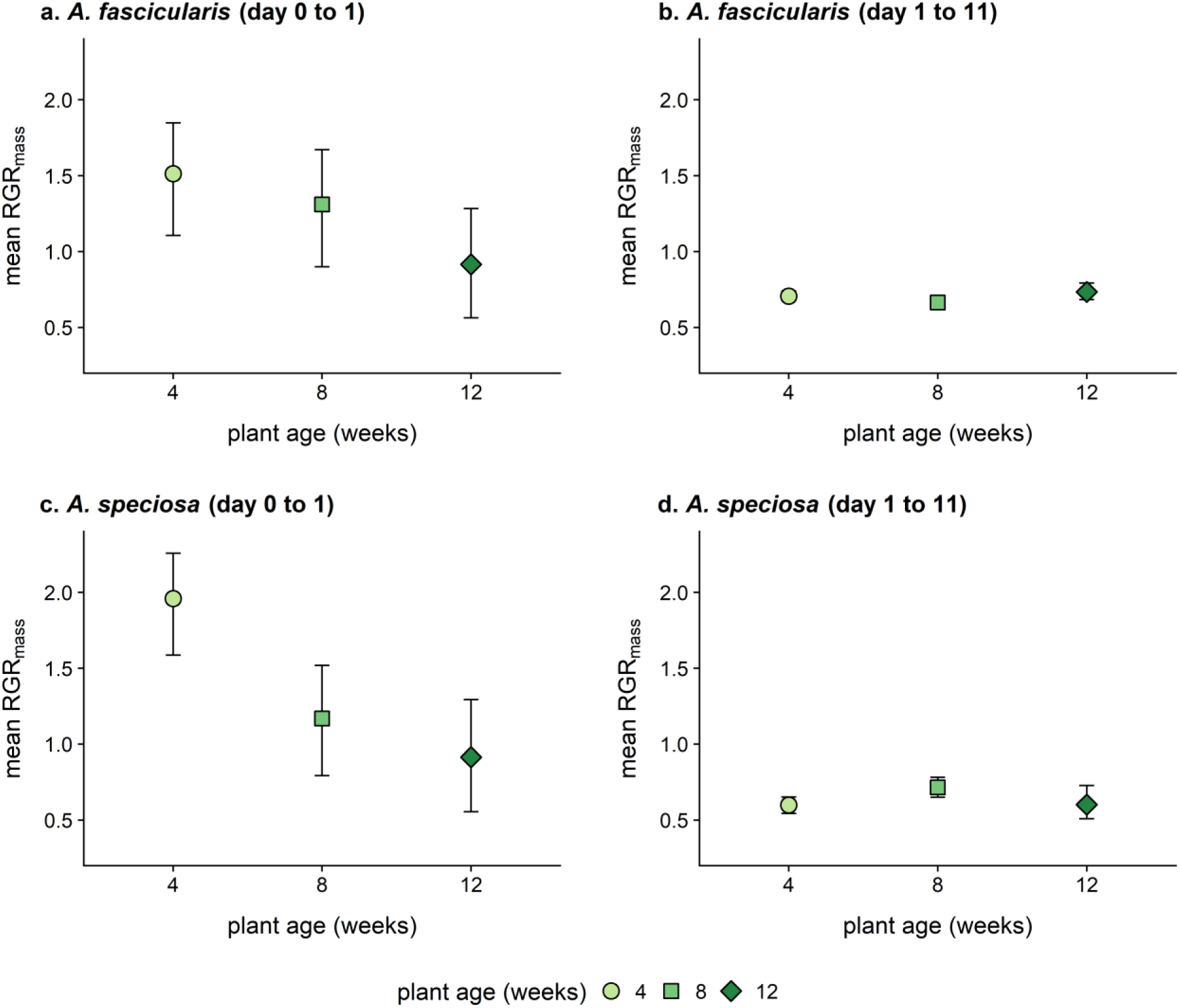
Interval-specific relative growth rates for caterpillars developing on narrow-leaved milkweed during a) experimental days 0 to 1 and b) experimental days 1 to 11, and for caterpillars developing on showy milkweed during c) experimental days 0 to 1 and d) experimental days 1 to 11. These figures show that the persistent negative effects of plant age on caterpillar size shown in Figs 3 and 4 emerges from growth differences in the first 24h of larval development. Point color and point shape represent plant age. Error bars represent 95% confidence intervals.

**Figure 6.**
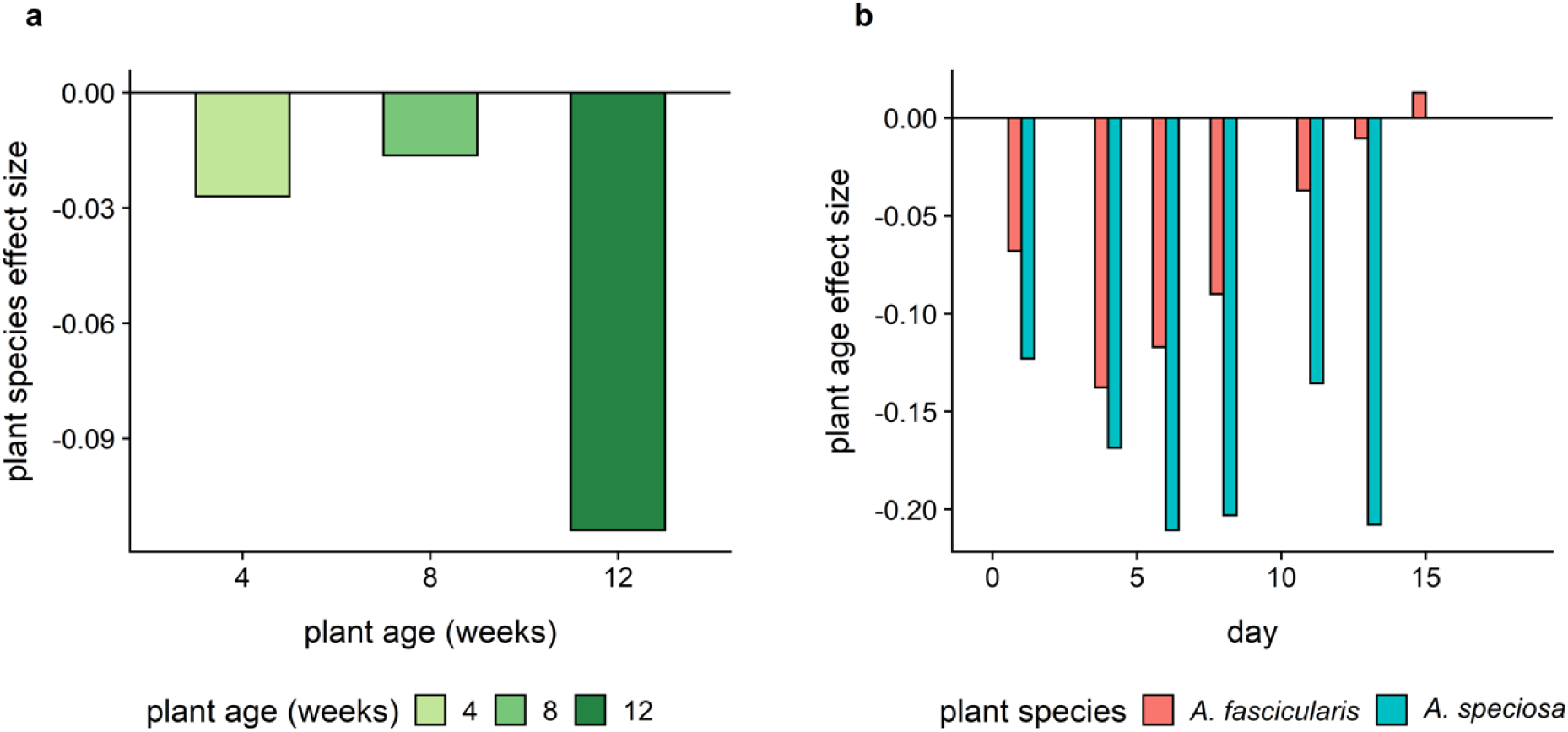
a) The mean plant species effect size for each plant age. These effect sizes represent the linear model coefficients for the effect of showy milkweed relative to narrow-leaved milkweed on surviving larval mass. Bar color represents plant age. Showy milkweed had a negative effect on larval mass in each plant age cohort, but this effect was larger in the oldest cohort. b) The mean plant age effect size for the surviving population on each experimental day, separated by host plant species. These effect sizes represent the linear model coefficient for plant age effects on surviving larval mass. Bar color represents plant species. The effects of plant age are consistently negative on showy milkweed. On narrow-leaved milkweed, the effect of plant age is generally negative, but the magnitude of these effects declines over time.

Caterpillars grew fastest on the youngest host plants in both species (Fig. 3-5, *plant age*: *F_2,72_*=9.6, *p*=0.0002). The overall relative growth rates of caterpillars were fastest on 4-week-old plants (0.82 mg/mg/day), and declined consistently on older host plants (8-week-old plants, 0.75 mg/mg/day; 12-week-old plants, 0.70 mg/mg/day, Fig. 4, see also Fig. S2 to S5). These differences in larval growth rates were established early, with diverging trajectories for caterpillars on plants of different ages appearing after the first experimental day (Fig. 3 and 5). The effect of plant age on monarch growth rates was stronger in the first 24h of the experiment than in the subsequent 10 days (Fig. 5, *plant age × interval: χ^2^_9_*=6.7; *p*=0.0099, see also Fig. S6), though this short, transient period of increased growth created persistent differences in caterpillar size throughout development (Fig. 3). Relative growth rates on 4-week-old plants were 1.9 times greater than those on 12-week-old plants across both plant species when looking at the interval from day 0 to day 1 (*plant age*: *F_1,96_*=17.2, *p*<0.0001, Fig. 5), and plant species identity did not have a significant effect on these growth rates (*plant species*: *F_1,96_*=0.4, *p*=0.53, Fig. 5). In contrast, in the interval from day 1 to day 11, caterpillars growth rates did not differ significantly among host plants of different ages (*plant age*: *F_1,38_*=0.58, *p*=0.45, Fig. 5), but did grow 9.1% faster on narrow-leaved milkweed compared with showy milkweed (*plant species*: *F_1,38_*=4.1, *p*=0.051, Fig. 5).

The effects of plant age on the realized growth rates of surviving larvae changed over the course of the experiment, as caterpillars died or left their host plant due to insufficient resources. The effects of plant age on caterpillar growth rates were variable but consistently negative throughout the experiment for showy milkweed, but these effects showed larger changes for caterpillars feeding on narrow-leaved milkweed (Fig. 6b). On narrow-leaved milkweed, the magnitude of the negative plant age effect declined throughout the experiment, and the few *(N*=4) caterpillars that survived to experimental day 15 showed a positive effect of plant age on larval growth rate (Fig 6b). This result suggests that while monarch caterpillars initially grew faster on younger plants, continued growth throughout the experiment was increasingly limited by host plant size.

### Analyses of maximum larval size

The expected maximum larval size attained, integrating both larval survival and growth, was greatest for caterpillars developing on larger, older plants across both host plant species (263 mg on 4-week-old plants, 317 mg on 8-week-old plants, 578 mg on 12-week old plants, *plant age: F_1,103_*=3.0, *p*=0.053, Fig. 7).

**Figure 7.**
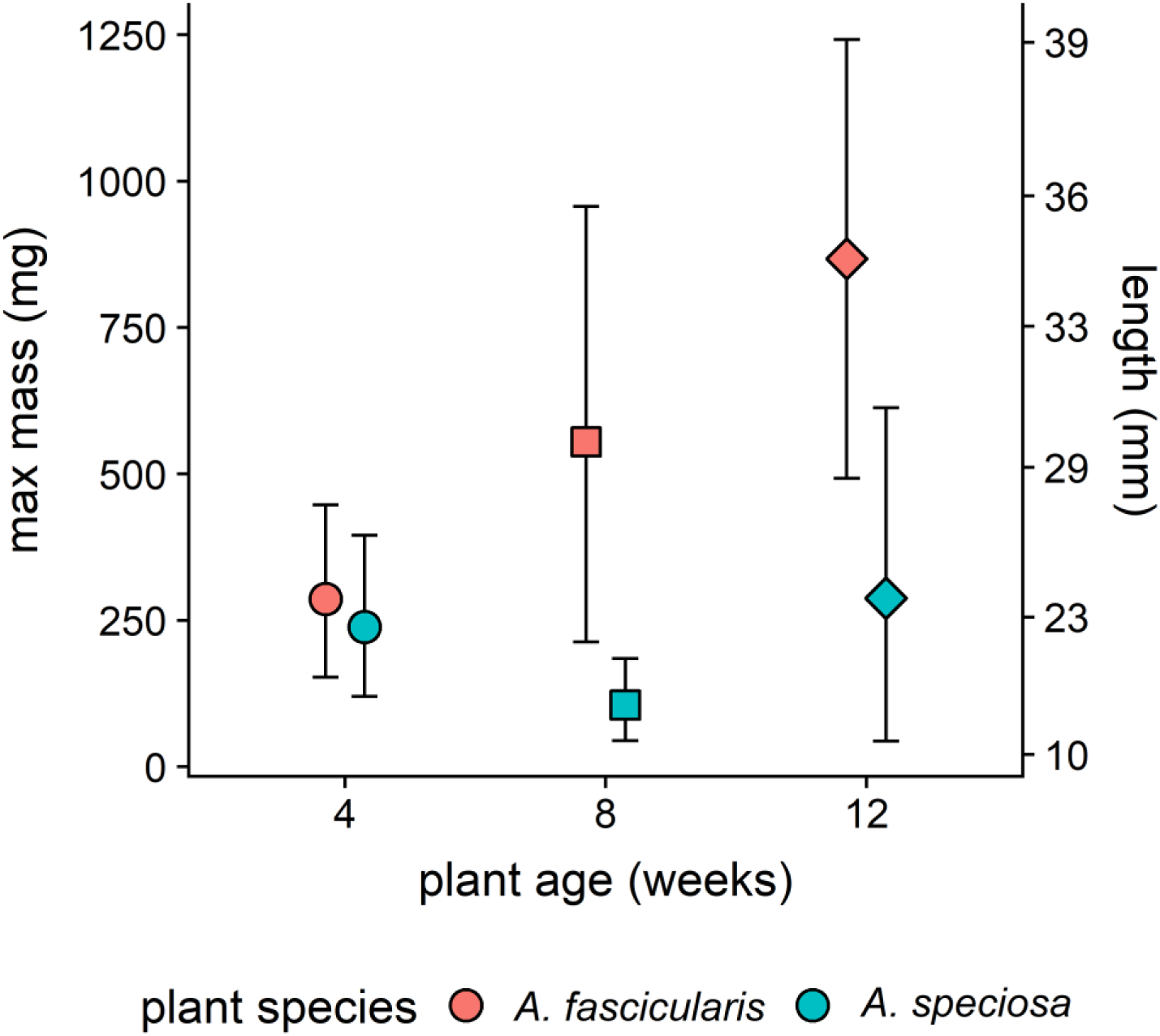
The maximum size (mass and length) attained by caterpillars developing on two host plant species of different ages. Color represents plant species, and point shape represents plant age. Error bars represent 95% confidence intervals.

Caterpillars also attained larger sizes growing on narrow-leaved milkweed than on showy milkweed. Across all cohorts, monarch larvae attained masses 2.7 times larger on narrow-leaved milkweed compared with showy milkweed (570 mg vs. 210 mg; *plant species*: *F_1,102_*=10.2, *p*=0.0018, Fig. 7). The difference between the maximum larval sizes attained on the two host plant species increased with plant age, from a 1.2-fold mean difference for 4-week-old plants to a 3-fold mean difference in 12-week-old plants, though these responses were variable and not statistically significant (*plant species × plant age: F_1,101_*=77.6, *p*=0.13). Comparable proportions of observed variation in maximum larval size were explained by plant species (*ΔR^2^*=0.087) and plant age (*ΔR^2^*=0.052).

### Analyses of plant damage

Caterpillars feeding on the youngest plants consumed a large proportion of available leaf area before leaving their host plant (Fig. 8a and 8b, *plant age: F_1,103_*=3.4, *p*=0.038), and caterpillars that stayed on the youngest host plants longer consumed nearly all available leaf material (Fig. 8c and 8d). The effect of plant age was particularly evident on showy milkweed; caterpillars left 4-week-old showy milkweed after consuming 26.1% of available leaf area, while caterpillars left 12-week-old showy milkweed after consuming only 5.6% of leaf area (Fig. 8b). Across all plant ages, percent damage was 1.4 times greater in narrow-leaved milkweed compared with showy milkweed (*plant species: F_1,102_*=1.4, *p*=0.24), and older showy milkweed deterred herbivory more strongly than younger plants. Among 4-week-old plants, the percent damage was 1.2 times higher in showy milkweed compared with narrow-leaved milkweed, but this pattern reversed in 8- and 12-week-old plants (2 times more herbivory in narrow-leaved milkweed among 8-week-old plants, and 2.5 times more herbivory in narrow-leaved milkweed among 12-week-old plants, *plant species × plant age: F_2,101_*=1.2, *p*=0.30).

**Figure 8.**
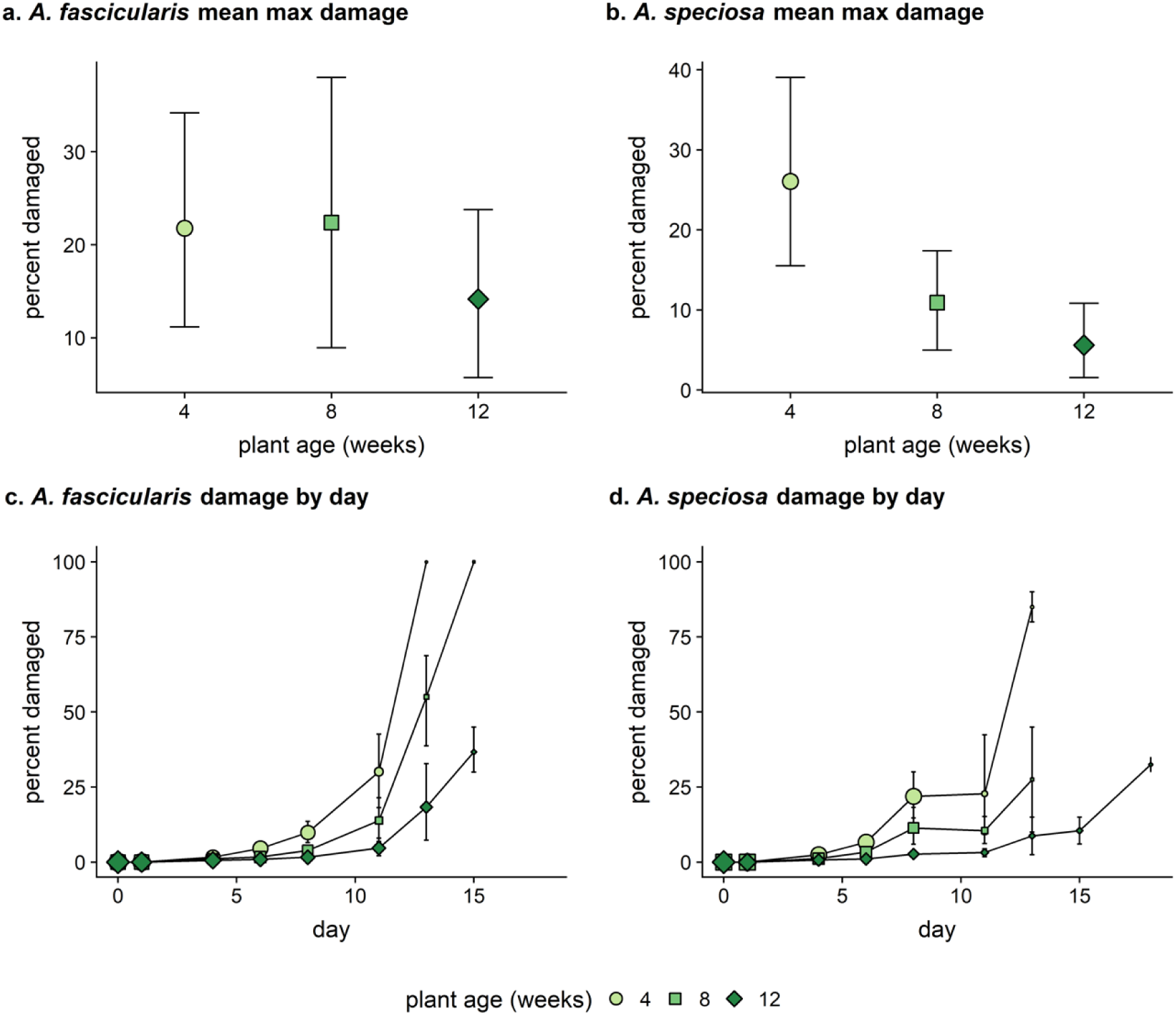
Plant damage by host plant species and age. Mean maximum herbivore damage for plants of each age cohort for a) narrow-leaved milkweed and b) showy milkweed. Mean maximum damage represents the percent of leaf area consumed by monarchs before leaving their host plant. Point color and point shape represent plant age. Error bars represent 95% confidence intervals. b) Percent damage on plants over time, showing the subset of plants with surviving caterpillars present at each time point. Point color and point shape represent plant age. Point size reflects the size of the surviving caterpillar population. Error bars represent 95% confidence intervals.

## Discussion

Taken together, these results show that species-specific and age-varying host plant traits significantly affect the growth and survival of monarch caterpillars. The plant traits that herbivores experience changed significantly over seasonal time following species-specific trajectories, and those changes in plant traits had strong effects on the developmental success of monarch larvae. Potentially in combination with seasonal changes in abiotic conditions and the biotic natural enemy community, these species-specific and age-varying changes in plant traits are likely to be important factors structuring seasonal windows of opportunity for monarch development.

Plant traits showed consistent differences between species and were strongly structured by plant age (Fig. 1). The species-specific differences between host plants increased with plant age for total stem length (Fig. 1a) and total number of leaves (Fig. 1b), reflecting species-specific differences in plant architecture. Because the growth form of showy milkweed is largely single-stemmed with large leaves, whereas narrow-leaved milkweed is generally branching with smaller leaves, total stem cross-sectional area and total leaf area are probably more indicative of the plant biomass available to herbivores than total stem length and total number of leaves. By comparison, total stem cross-sectional area (Fig. 1c) and total leaf area (Fig. 1d) showed relatively non-diverging ontogenetic trajectories suggesting that, despite large differences in their architecture, the plant biomass available to herbivores did not diverge between species as markedly over ontogeny as other species-specific traits, including defensive traits (Fig. 1e and 1f). Broadly, these results indicate that the traits experienced by herbivores are strongly influenced by plant age. While host plant species identity was also informative, plant age often explained a comparable proportion of the observed variation in plant traits in this study. These findings extend the meta-analytic dataset described by Barton and Koricheva (2010) which documented generally increasing constitutive chemical defenses from the seedling stage to maturity in herbaceous plants, but lacked a sufficient sample size of studies to identify general ontogenetic patterns in physical defense traits with herbaceous plants (but see Traw and Feeny 2008). The results of this current study show significant changes in both plant defense traits over ontogeny, but the trajectories of these traits also differed strongly between the two milkweed species.

By comparison, plant age explained substantially more variation in overall larval growth rate than plant species (Fig. 4). Across larval development, monarch caterpillars grew fastest on the youngest plants of both species, and this overall pattern was strongly (and unexpectedly) driven by large differences in growth rate during the first 24h of larval development (Fig. 5). Plant age-associated differences in larval growth rate during the first day after egg introduction created substantial differences in larval size that persisted throughout the rest of larval development (Fig. 3). This result is consistent with a previous study showing that monarch caterpillars grew faster on milkweed leaves with partially severed petioles (and thus reduced latex pressure) during the first 2-4 days of larval development on four out of nine species of milkweed examined (Zalucki et al. 2001); in both studies, early instar caterpillars grew faster on host leaves with reduced latex exposure. These findings are also consistent with studies indicating that adult monarchs preferentially oviposit on younger host plants (Zalucki and Kitching 1982), as well as the recent vegetative regrowth of host plants that have been strategically mowed for habitat management (Fischer 2015, Haan and Landis 2019, Knight et al. 2019). Similar preferential herbivory on rapid regrowth has been observed in other systems in response natural disturbance regimes (e.g., Spiller and Agrawal 2003). Our results suggest that plant age is a key determinant of variation in this defensive trait, and show that the strongest effects of these age-associated differences in plant traits on growth rate occur in the first 24h of larval development.

Monarch caterpillars experienced greater developmental success (i.e., attained larger maximum larval sizes) on narrow-leaved milkweed than on showy milkweed (Fig. 7), and the difference between host plant species was particularly strong for older host plants (Fig. 7). These findings are consistent with our observation that older plants showed more strongly differentiated species-specific plant traits in this study, while younger plants of both species were unexpectedly similar in their traits. These two milkweed species express distinct plant defense syndromes as mature plants (Agrawal and Fishbein 2006). In our study, species-level differences emerged over ontogeny as the defensive traits of these species diverged with increasing plant age (Fig. 1e and 1f). On the oldest host plants, both growth (Fig. 4 and 6a) and survivorship (Fig 2c) were strongly structured by plant species; on both counts, caterpillars developing on narrow-leaved milkweed fared better than those developing on showy milkweed. These patterns are consistent with the different seasonal windows of opportunity that have been previously observed for monarchs feeding on these two host plants (Yang and Cenzer 2019): while monarchs showed two seasonal windows of opportunity on narrow-leaved milkweed, those feeding on showy milkweed only showed the early season window. We suggest that increasing plant defense traits over ontogeny could limit late season windows of opportunity in showy milkweed. The findings of our current study are also consistent with the hypothesis that the two seasonal window of opportunity observed on narrow-leaved milkweed could correspond to a “double-dipping” herbivore strategy (*sensu* White 2015, Che-Castaldo et al. 2019) in which monarch larvae successfully use both vigorously growing and senescing plant tissues. Our findings indicate that the bounds of the early season window of opportunity may be influenced by temporally varying resource quantity (*i.e.*, plant size) and quality (as affected by age-varying defensive traits). Future studies will be necessary to more specifically examine how increasingly senescent plant traits affect larval success in the second window of opportunity observed in this system.

In this study, differences between plant species in the effect of plant age became more apparent as the experiment progressed (Fig. 6b); while caterpillars generally developed better on younger plants than on older plants, the effect of plant age was more consistently negative throughout monarch development on showy milkweed (Fig. 6b). In comparison, when developing on narrow-leaved milkweed, caterpillars early in the experiment showed a smaller negative effect of plant age relative to those developing on showy milkweed, and these negative effects became weaker throughout the experiment (Fig 6b). This plant age effect trajectory on narrow-leaved milkweed illustrates the multiple and potentially conflicting effects of plant age: while younger plants provided higher *quality* resources that allowed for faster larval growth rates initially, older plants provided greater resource *quantity* over a longer developmental timescale. These changes in the developmental limitations imposed by seasonally varying resource quality and quantity are further supported by observed patterns of herbivore damage and larval survival. On the youngest plants, the developmental success of larval monarchs appeared to be ultimately limited by the availability of host plant biomass (*i.e.,* resource quantity). Caterpillars on the youngest plants fed on less-defended (i.e., higher-quality) resources and grew fast (Figs. 1 and 4); they often consumed a substantial proportion of their host plants before starving or attempting to disperse to a second host plant (Fig. 8). As a result, these caterpillars showed steep and short survivorship curves on both host plant species; in general, these caterpillars grew fast and died young (Fig. 2). In comparison, caterpillars developing on the oldest host plants seemed to be limited by the *quality* of host plant biomass as constrained by plant defense traits. These caterpillars showed the slowest growth rates (Fig. 4), but rarely consumed their entire host plant (Fig. 8), and showed the longest survivorship curves (Fig. 2).

The relative importance of milkweed *quality* and *quantity* as factors that structure seasonal windows of opportunity for monarch development could also depend on the density of milkweeds in available habitat patches. This experiment was conducted with singular host plants as replicates, where attempted dispersal by larvae below the pupation threshold size was assumed to be fatal. This assumption is likely to be a reasonable one in habitats where individual plants are widely spaced, where biotic or abiotic conditions limit the ability of monarch caterpillars to move between neighboring plants (e.g., due to increased thermal stresses or predation risk), or if monarchs show limited abilities to locate second host plants. Alternatively, high-density patches of young milkweed plants could potentially provide high-quality host plant resources with reduced plant-quantity constraints; this suggests that higher density patches could potentially allow for earlier seasonal windows of opportunity, consistent with the results of previous field experiments (Yang and Cenzer 2019). Further studies specifically examining the context-dependent risk of plant-to-plant movement in milkweed patches of varying density, in different habitats, at different larval sizes, and at different times of year would be valuable to better understand how plant density could affect seasonal window of opportunity for monarch development. Moreover, while this study investigated the effects of plants traits in two milkweed species during their first growing season, additional studies assessing other host plant species, additional plant traits (including physical, chemical and indirect defense traits), and a wider range of plant ages (especially considering plants in their second growing season and beyond) will be necessary to assess the generality of the patterns observed here.

The results of this study indicate that age-varying plant traits likely play a strong role in structuring seasonal windows of opportunity for monarch caterpillars. However, because this experiment was designed to isolate the effects of age-varying plant traits without the contributing effects of temporally variable abiotic and top-down factors, the role of seasonal variation in climatic conditions and natural enemy interactions remains uncertain. Both additional factors are likely to interact with the effects of plant trait variation in nature; for example, delayed growth rates on lower quality host plants could expose larvae to greater predation risk. While both additional factors are likely to affect seasonal windows of opportunity for larval development, either independently or interactively with seasonal changes in host plant quality or quantity, separate experimental studies will be necessary to quantify their effects.

More broadly, these findings contribute to the general observation that temporal variation in plant traits can strongly affect plant-herbivore interactions (e.g., Van der Wal et al. 2000, Van Asch et al. 2007, Barton and Koricheva 2010, Che-Castaldo et al. 2019). The results of this study indicate that the effects of plant age on monarch developmental success are comparable to and sometimes larger than those of plant species identity. Acknowledging substantial temporal variation in plant traits does not diminish the importance of species-level trait assessments; expectations about how plant traits affect herbivores are often usefully structured around species-level characterizations, and such studies can identify clusters of species that share key traits (Agrawal and Fishbein 2006). In combination with such species-level trait assessments, the temporal dimensions of plant age and seasonal variation provide additional orthogonal axes to examine variation in plant defense traits and their effects on herbivores.

These results may also suggest some specific implications for our understanding of milkweed-monarch interactions in a changing world, and the potential for milkweed limitation in the population dynamics of monarchs (Nail et al. 2015, Pleasants et al. 2016, Inamine et al. 2016, Thogmartin et al. 2017), and especially in western North America (Espeset et al. 2016, Pelton et al. 2019). If age-varying plant traits have strong effects on the developmental prospects of monarchs generally, monarchs may experience changing constraints on larval development as their host plant traits develop through the season. In particular, the development of monarch larvae in the early season could potentially be limited by small host plant size, even in habitats with abundant host plant resources later in the season. Conversely, if later-season milkweeds generally present stronger defensive traits than early-season plants, monarchs could potentially experience reduced growth rates during periods of lower resource quality even when the apparent availability of host plant resources is high. Because these potential seasonal limitations are mediated by changes in resource quality as much as resource quantity, estimates of milkweed abundance and spatial distribution by themselves may not capture a key temporal dimension of the dynamic resource landscape. If a wider range of milkweed species show the kinds of species-specific and age-varying traits observed in this current study, it would suggest that migrating monarchs face a complex and dynamic landscape of potential host plants with traits that are affected by phenology and ontogeny as much as species distributions. The complexity of this dynamic resource landscape likely presents a challenge for migrating monarchs as well as the ecologists that aim to study them. Developing a more temporally explicit approach may be necessary to assess the combined effects of plant age and species identity on the spatial distribution and temporal availability of milkweed resources on a continental scale. Further, it is unclear how monarch migrations and the dynamics of this seasonally variable landscape will change with global warming. The age of host plants that migrating monarchs encounter each year is likely to be affected by both the environmental cues that influence milkweed phenology, as well as the continental-scale drivers of monarch migration. The potential for significant mis-matches in the relative phenologies of milkweeds and monarchs remains uncertain, though the magnitude of observed plant-age effects in this study suggests that the consequences of such phenological mis-matches, if realized, could be substantial. Further studies will be necessary to identify the environmental cues that drive phenological responses in a range of milkweed species, and how phenological variation across different species distributions affects the overall spatiotemporal availability of milkweed resources throughout each season.

## Acknowledgements

We thank August Higgins, Jennifer McKenzie, Jessica Aguilar and Nicholas Rasmussen for their assistance. This study was supported by a National Science Foundation CAREER award to LHY (DEB-1253101).

## Appendix S1

**Figure S1.**
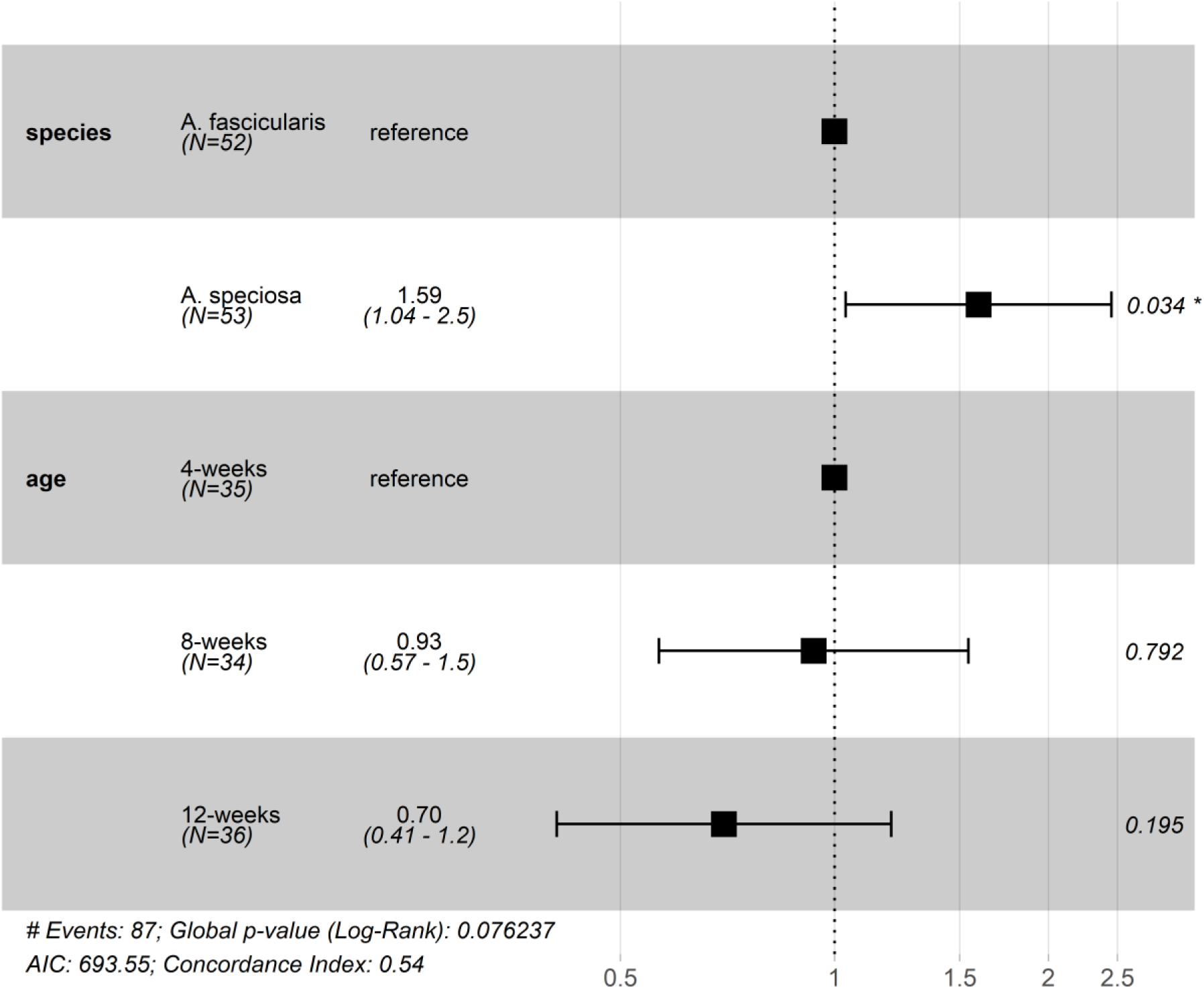
Hazard ratios from a Cox proportional hazard (survivorship) model with plant species and plant age as explanatory factors. The first column indicates the explanatory factor, the second column indicates the levels of each factor, the third column represents the estimated hazard ratio with 95% confidence intervals in parentheses, the fourth column shows the estimated hazard ratio and confidence intervals graphically, and the fifth column shows the *p*-value for each non-reference factor level.

**Figure S2.**
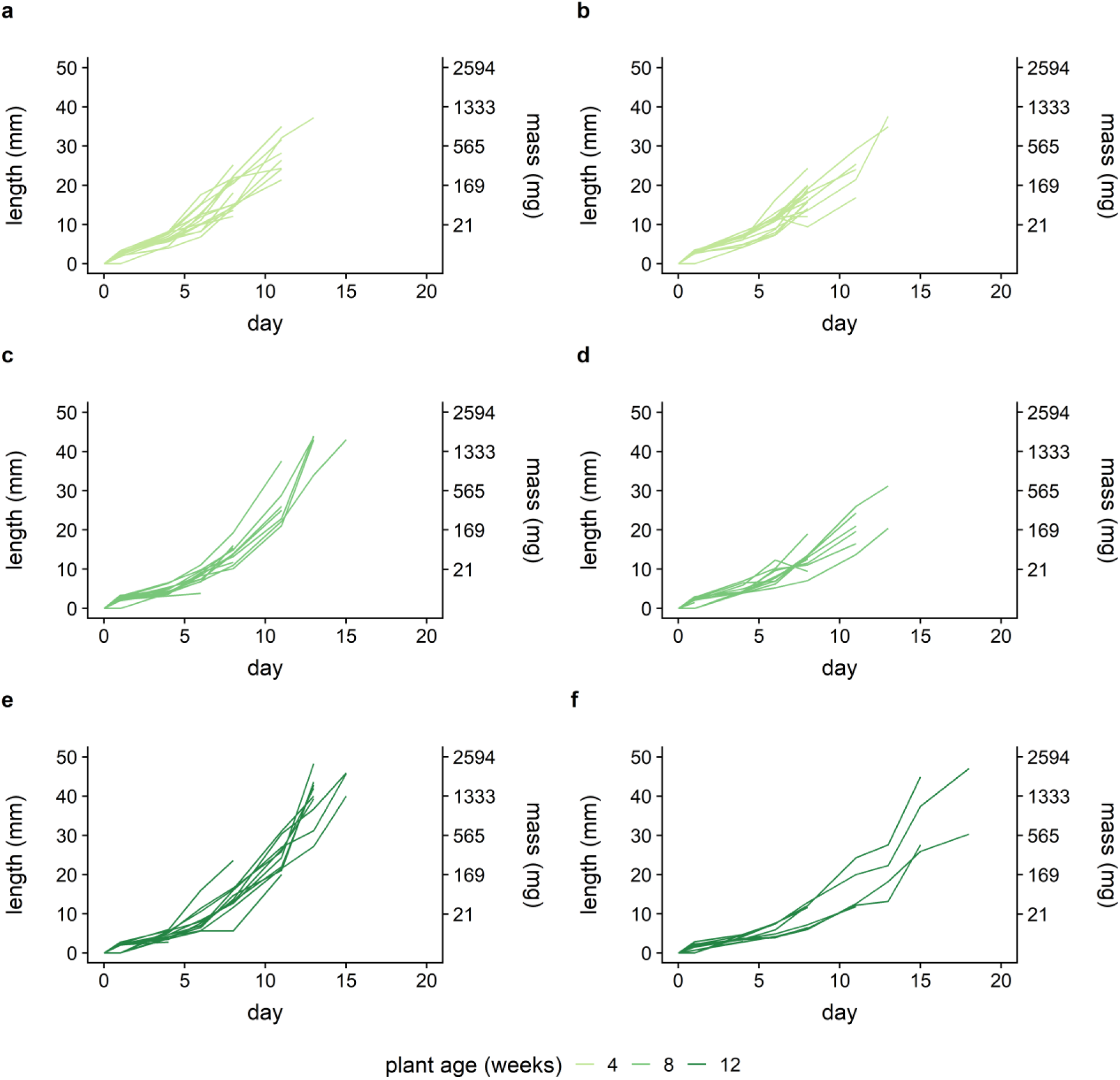
Individual growth trajectories of caterpillar size in each plant species and age cohort; each line presents the size of an individual caterpillar. Panels in the left column (a, c and d) present data from caterpillars reared on narrow-leaved milkweed, while panels on the right column (b, d and f) present data from caterpillars reared on showy milkweed. Line color represents plant age. Compared with Fig. 3, these data are shown on an untransformed axis, and with length on the primary axis and mass on the secondary axis, in order to present a minimally processed overview of individual caterpillar growth.

**Figure S3.**
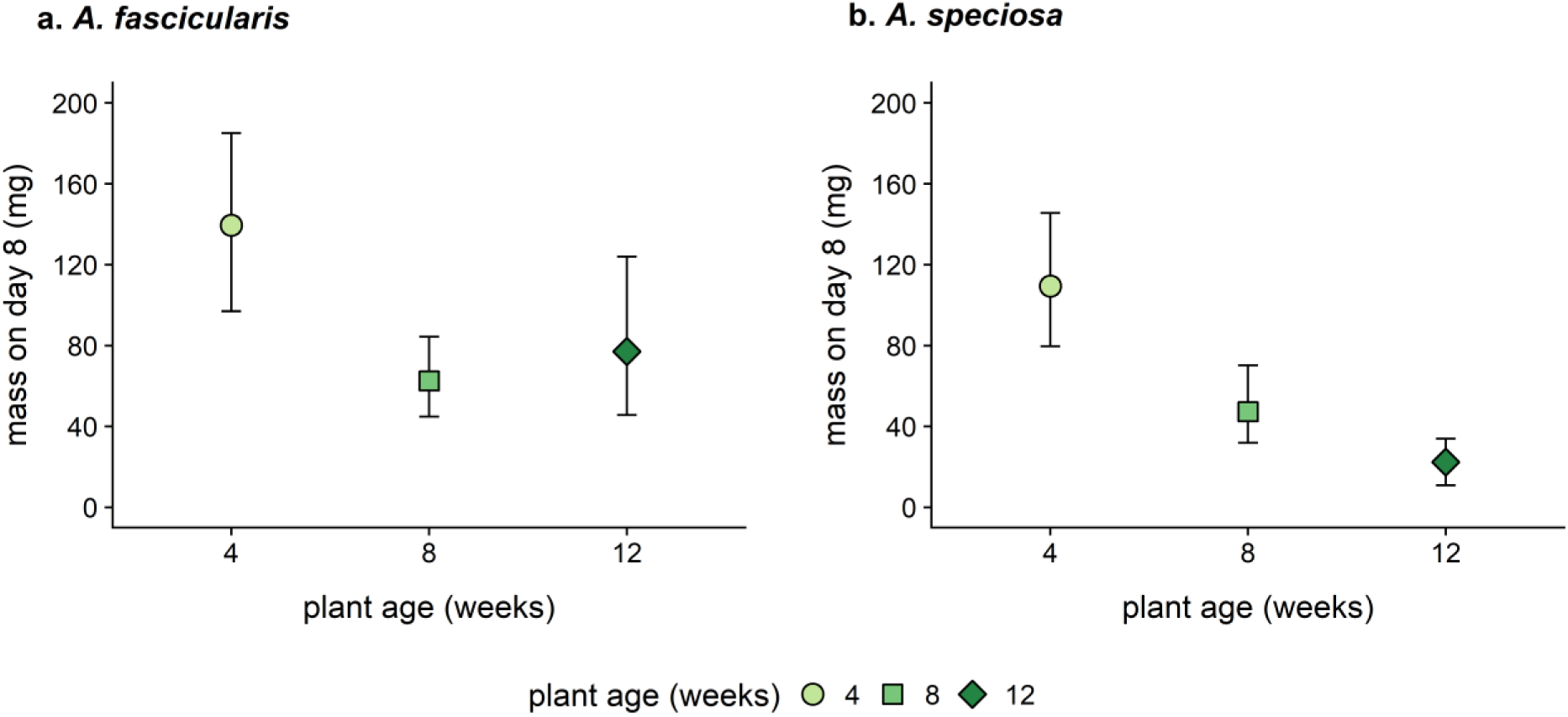
Mean mass on day 8 for caterpillars that developed on each plant species at each plant age. This metric is similar to the overall relative growth rate calculated via log-linear regression and shown in Fig. 4 because both metrics assess growth from the beginning of the experiment, when initial sizes were zero, and both represent a similar overall sample size (*N*=71 in this figure vs *N*=74 in Fig. 4). Point color and shape represent plant age. Error bars represent 95% confidence intervals.

**Figure S4.**
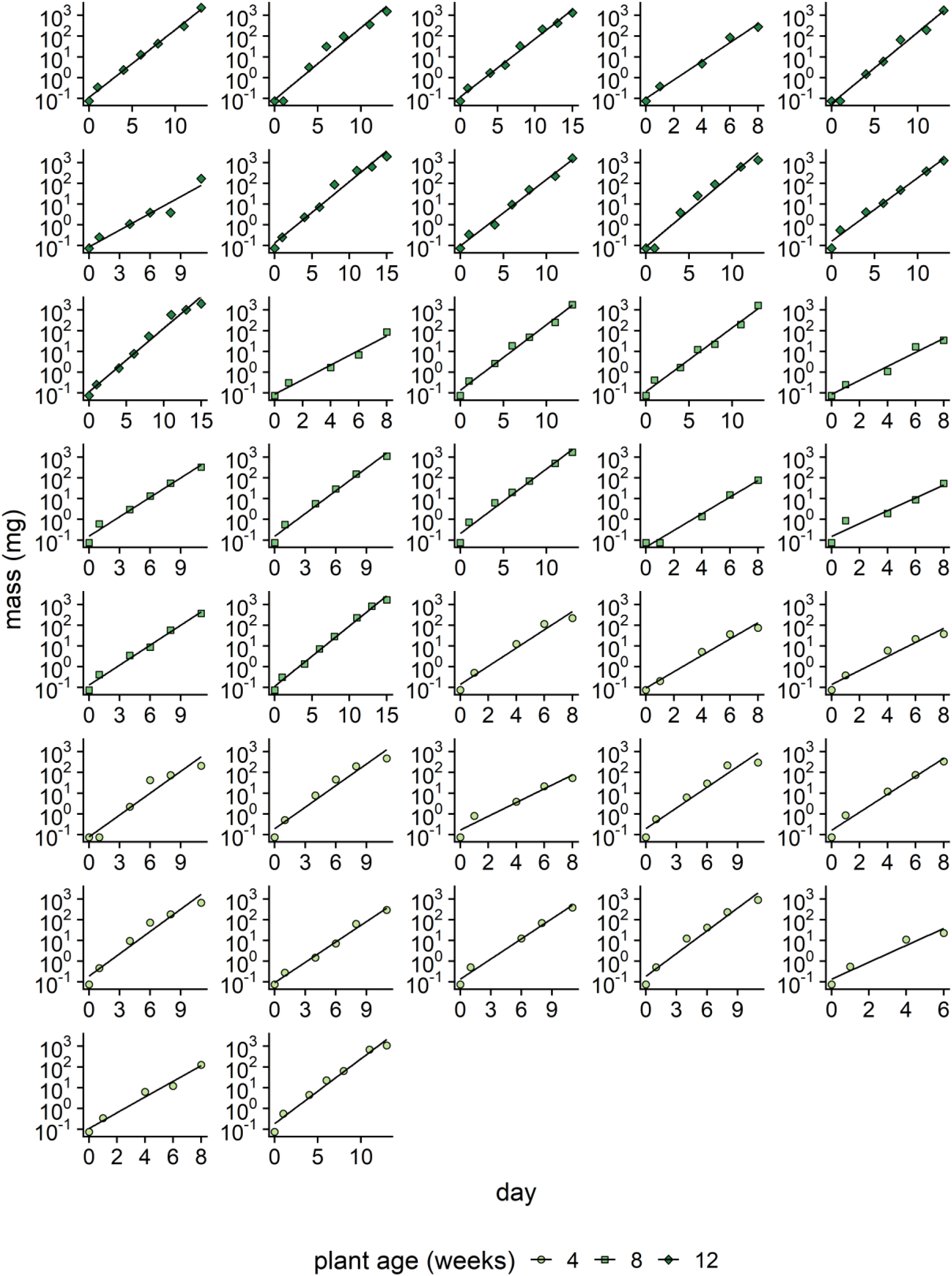
Individual log-linear fits for caterpillars reared on narrow-leaved milkweed. Point color and shape represents plant age, and the black line represents the best fit log-linear regression.

**Figure S5.**
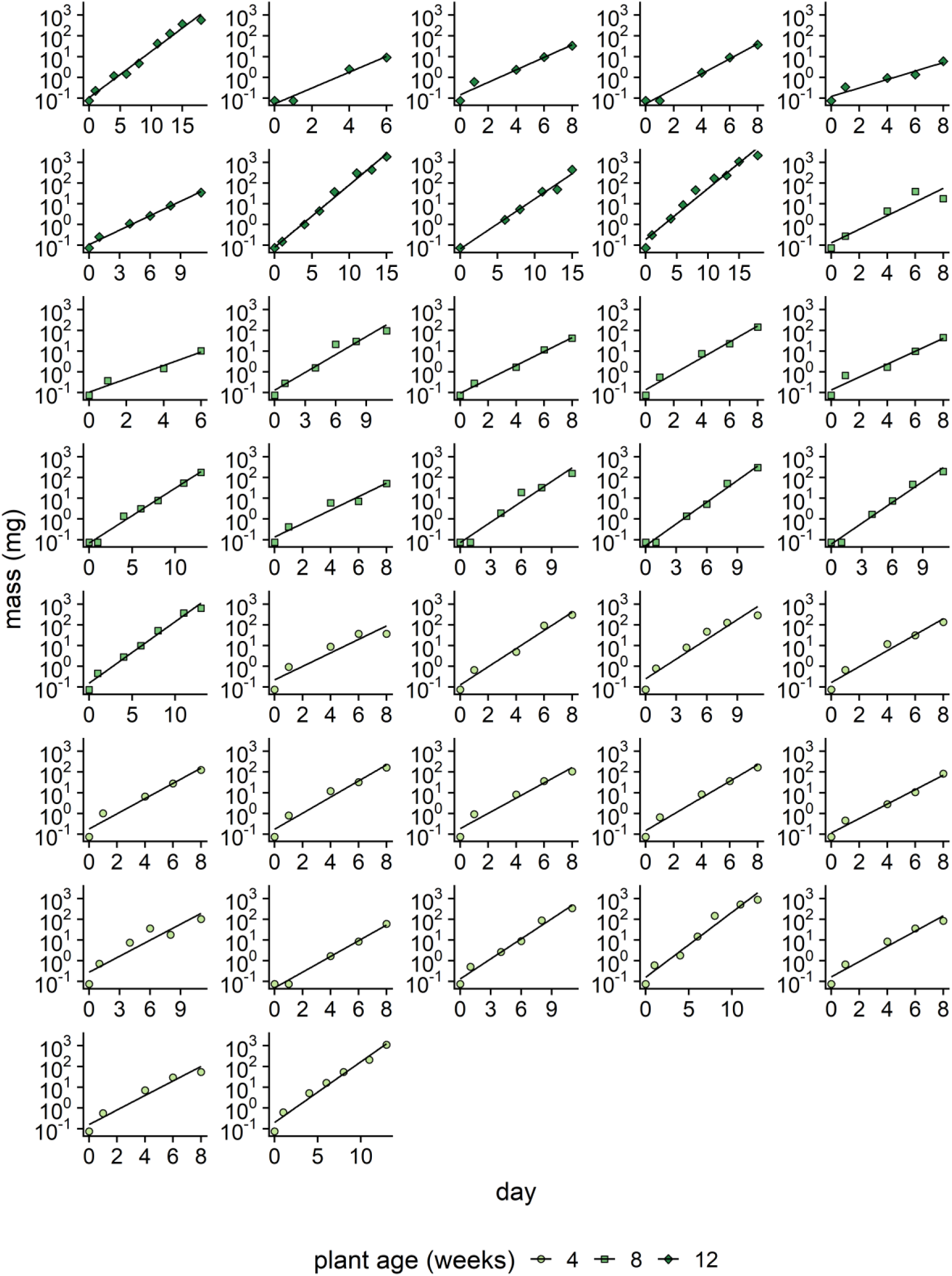
Individual log-linear fits for caterpillars reared on showy milkweed. Point color and shape represents plant age, and the black line represents the best fit log-linear regression.

**Figure S6.**
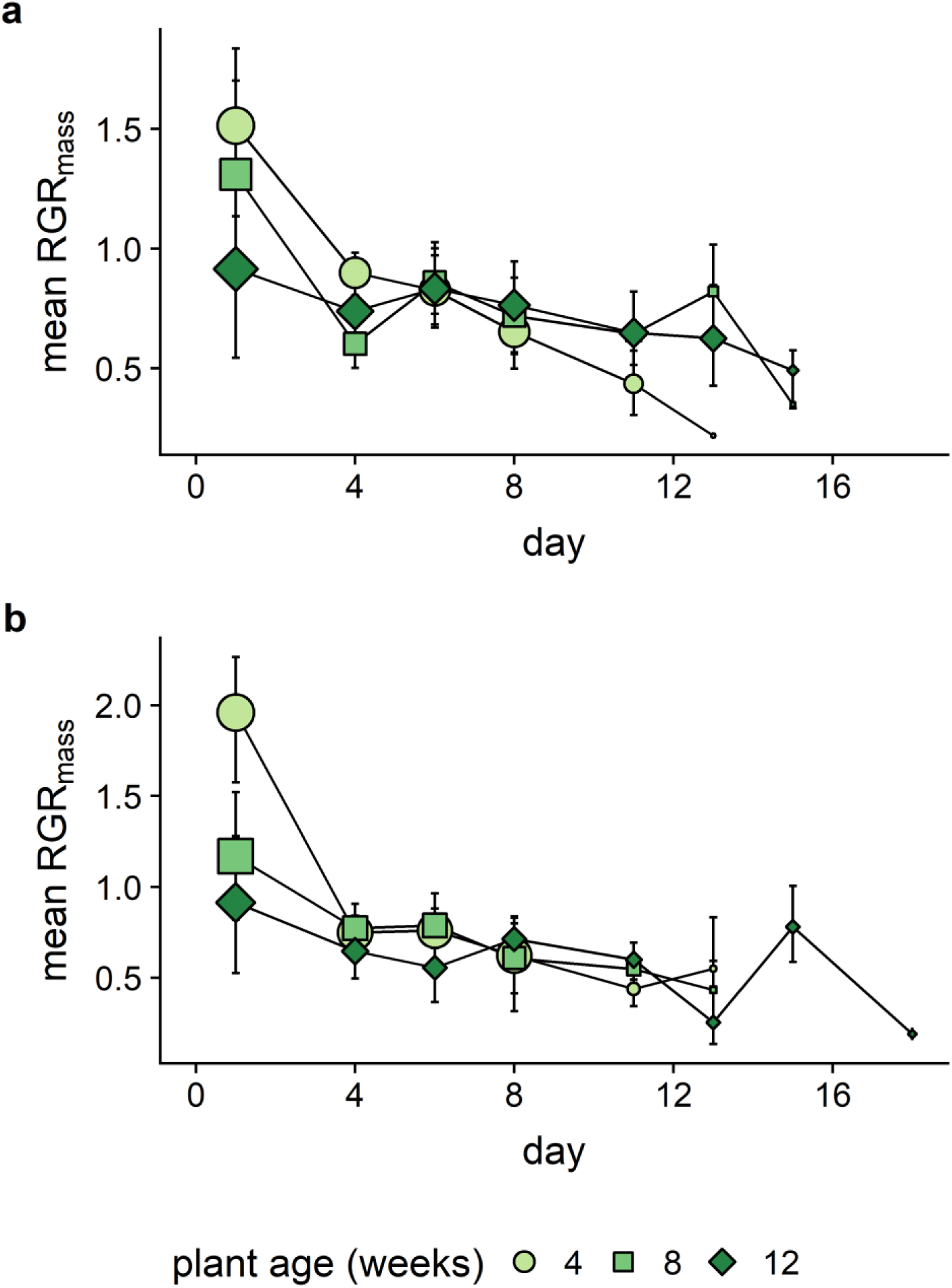
Interval-specific relative growth rates calculated for all adjacent observations on a) narrow-leaved milkweed and b) showy milkweed. This figure is similar to Fig. 5, but presents growth data for all possible adjacent intervals. Point size reflects the size of the surviving population. Point color and shape represent plant age. Error bars represent 95% confidence intervals.

